## Supplemental Information for "Structure and dynamics of endogenous protein complexes in human heart tissue captured by native nanoproteomics"

**Table of Contents**

**Supplementary Methods**

Synthesis of iron-oleate precursor…..…………………………………….……..……….S-4-5

Synthesis of iron-oleate precursor….…………………………………….……..…………S-5

Synthesis of 8 nm magnetite ($Fe_3O_4$) nanocrystals ….……….……….……..……..…S-5-6

Synthesis of $Fe_3O_4$-BAPTES NPs (NP-BAPTES)….……….……….……..…….…….S-6

Synthesis of $Fe_3O_4$-BAPTES-Peptide NPs (NP-Pep)….…………..……..……..…….S-6-7

Top-down RPLC-MS/MS Analysis of cTnI, cTnT, and TnC monomers..………..….S-7-8

Size exclusion chromatography (SEC) for online buffer exchange (OBE) and native MS analysis of the cTn(I-T-C) complex.…………………………………….……..………..S-8-9

**Supplementary Tables**

Table S1. Available clinical data for non-failing donor hearts used……………….......S-10

Table S2. cTn proteoforms identified……………………………………..…....…..…S-11

Table S3. Summary of native TIMS-MS CCS ($Å^2$) values for TnC monomer, cTn(I-C) dimer, and cTn(I-T-C) heterotrimer complex. Theoretical CCS values were determined using the IMPACT method………………………………..……….…………..…S-12

Table S4. Summary of native TIMS-MS CCS ($Å^2$) values for the cTn(I-T-C) complex incubated with 0, 25, 50, and 100 mM EGTA..............................................……S-13

**Supplementary Figures**

Supplementary Figure S1. Optimization of elution buffer solution for cTn enrichment with NP-Pep……………………………..……………….……..……………………..…....S-14

Supplementary Figure S2. High reproducibility of inter-batch enrichment by NP-Pep……...........................................................................................................…......S-15

Supplementary Figure S3. Evaluation of NP-Pep native enrichment performance using RPLC-MS/MS..….............................................................................................….…..S-16

Supplementary Figure S4. Reproducibility of the native NP-Pep enrichment capturing cTn monomers in human cardiac tissues using RPLC-MS/MS................…..…………….…S-17

Supplementary Figure S5. Native size exclusion chromatography for online buffer exchange (OBE) enables MS analysis of protein and protein complexes..........….……S-18

Supplementary Figure S6. Reproducibility of native cTn enrichment and SEC-OBE native MS analysis.............................................................................................................………S-19

Supplementary Figure S7. Isotopically resolved native MS spectra of endogenous cTn heterotrimer complex (~77 kDa)….......................................................................…..S-20

Supplementary Figure S8. Complex-up native MS analysis reveals cTn complex stoichiometry.…......................................................……………………….…………S-21

Supplementary Figure S9. Native top-down fragmentation map of cTnT monomer..………………………………………………….....................................S-22

Supplementary Figure S10. Native top-down fragmentation map of cTnI monomer..………………………..…………………………………………...……..…S-23

Supplementary Figure S11. Relative proportion of $Ca^{2+}$ bound in each TnC binding domain.......................................................................................................................S-24

Supplementary Figure S12. Localization of calcium binding in TnC monomer site III by native top-down MS (nTDMS)..........................................................................S-25

Supplementary Figure S13. Localization of calcium binding in TnC monomer site IV by native top-down MS (nTDMS)........................................................................................S-26
Supplementary Figure S14. Localization of calcium binding in TnC monomer site II by native top-down MS (nTDMS)........................................................................................S-27
Supplementary Figure S15. Optimization of TIMS parameters for high resolution native MS analysis of bovine serum albumin (BSA)...............................................................S-28
Supplementary Figure S16. Plot of CCS value as a function of charge state by native TIMS-MS..................................................................................................................S-29

**Supplementary Methods**

**Synthesis of N-(3-(triethoxysilyl)propyl)buta-2,3-dienamide.** 3-Butynoic acid (2.10 g, 1 eq., 25 mmol), 2-chloro-1-methylpyridinium iodide (7.00 g, 1.1 eq., 27 mmol), and dichloromethane (250 mL) were added to a round bottom flask. Next, a solution of (3-aminopropyl)triethoxysilane (5.53 g, 1 eq., 25 mmol), N,N-diisopropylethylamine (6.46 g, 2 eq., 50 mmol), and dichloromethane (125 mL) was prepared separately and added to the stirring round bottom flask. The reaction mixture was allowed to reflux for 1 hour. The crude product was concentrated *in vacuo* and partially redispersed in ethyl acetate, allowing for the separation of BAPTES from the insoluble 2-chloro-1-methylpyridinium iodide by gravity filtration. The product was concentrated *in vacuo* and purified twice with column chromatography using a gradient of 50:50 to 100:00 ethyl acetate: hexane to yield BAPTES and its alkyne isomer as a clear, orange oil (4.80 g, 67% yield). To effectively isomerize the propargylic isomer to N-(3-(triethoxysilyl)propyl)buta-2,3-dienamide, anhydrous acetonitrile (120 mL), BAPTES isomers (3.38 g, 1 eq., 11.78 mmol), and tribasic potassium phosphate (2.50 g, 1 eq. 11.78 mmol) were added to a round bottom flask. The reaction was allowed to stir at room temperature for 2 hours before gravity filtration and subsequent concentration *in vacuo*. The product was purified with flash column chromatography using a gradient of 70:30 to 100:00 ethyl acetate: hexane to yield a clear, yellow oil. The final product was concentrated *in vacuo* for further use; $^{1}$H NMR (500 MHz, Chloroform-*d*) δ (ppm) 5.99 (s, 1H), 5.62 (t, $J$ = 6.6 Hz, 1H), 5.20 (d, $J$ = 6.7 Hz, 2H), 3.83 (q, $J$ = 7.0 Hz, 6H), 3.31 (q, $J$ = 6.7 Hz, 2H), 1.66 (m, 2H), 1.24 (t, $J$ = 7.0 Hz, 9H), 0.66 (m, 2H); $^{13}$C NMR (125 MHz, Chloroform-*d*) δ (ppm) 211.56, 164.33, 91.02, 80.33, 58.47, 42.02, 23.02, 18.30, 7.66; ESI-MS for $C_{13}H_{25}NO_4Si$ $[M+H]^+$ observed: 288.162 *m/z*, $[M+H]^+$ calculated: 288.162 *m/z*. (2.30 g, 68% yield).

**Synthesis of iron-oleate precursor.** Iron oleate was synthesized using a previously established method[26]. In a typical synthesis, iron (III) chloride hexahydrate (10.8 g, 40 mmol) was first dissolved in a mixture of 80 mL ethanol and 60 mL nanopure water in a three-neck round bottom flask (500 mL) containing a Teflon-coated egg-shaped (1-1/4" x 5/8") magnetic stir bar. Sodium oleate (36.5 g, 120 mmol) was then quickly added to the iron chloride solution along with 140 mL n-hexane. The resulting solution was then allowed to stir until the sodium oleate was completely dissolved. Afterwards, the reaction solution was heated to 70 °C for a 4-h reflux under a $N_2$ blanket. Upon completion, the reaction solution was cooled to room temperature, and the upper organic layer containing the iron oleate was washed three times with 30 mL nanopure water in a 250 mL separatory funnel. After washing, the iron oleate was concentrated *in vacuo*. Finally, the resulting iron oleate was transferred into a 100 mL round bottom flask, connected to a Schlenk line, and placed under vacuum overnight. For storage, the iron oleate was well sealed in a glass vial and placed in a desiccator.

**Synthesis of 8 nm magnetite ($Fe_3O_4$) nanocrystals.** Iron oleate precursor and magnetite nanoparticles were synthesized following published protocol, with minor modifications.[19] Briefly, the NPs were synthesized using 10 mmol (9.0 g) iron oleate, 5.5 mmol (1.56 g) oleic acid, and a 4:1 ODE:TDE (40 g: 10 g) solvent mixture. The mixture was stirred and degassed on a Schlenk line at 110 °C for 3 h, heated to 300 °C at a heating rate of 3.3 °C/min under a nitrogen flow, and then allowed to boil for 30 min. After three centrifugation wash cycles (10,000 × *g*, 20 min) using EtOH, the resulting NPs were then dried under vacuum, weighed, and redispersed in n-hexane at a concentration of 20 mg/mL for further use.

**Synthesis of $Fe_3O_4$-BAPTES NPs (NP-BAPTES).** In a typical optimized large-scale synthesis of silane functionalized NPs, $Fe_3O_4$ NPs (6 mL from a 20 mg/mL stock) were added to anhydrous n-hexane (300 mL) in a 500 mL round bottom flask equipped with a Teflon-coated egg-shaped magnetic stir bar (1–1/4" × 5") to achieve a total NP concentration of 0.4 mg/mL. After the reaction mixture was heated to 60 °C with stirring (900 rpm), BAPTES (1.65 mL) was added dropwise to the flask for a 0.55% (v/v) total concentration of trialkoxysilane reagent, followed by dropwise addition of a small amount of acetic acid (30 μL) for an acidic catalyst concentration of 0.01% (v/v). After reacting at 60 °C for 24 h, the precipitate was collected and washed one time with n-hexane, one time with n-hexane/acetonitrile (v/v, 4:1), and one more time with n-hexane via centrifugation (10,000 × *g*, 10 min) to remove excess silane molecules and surfactants. The NPs were then dried under vacuum for later use.

**Synthesis of $Fe_3O_4$-BAPTES-Peptide NPs (NP-Pep).** 10 mg of NP-BAPTES were added to a 4-dram vial and dispersed in 2 mL of acetonitrile. 10 mg of cTnI-binding peptide (HWQIAYNEHQWQC) were added to a separate 4-dram vial and dissolved in 8 mL of nanopure water. The pH of the peptide solution was adjusted to pH 8.0 by the addition of 75 µL of 1.0 M ammonium carbonate buffer pH 9.0 during simultaneous water bath sonication. The pH-adjusted peptide solution was added into the 4-dram vial containing the NP dispersion under water bath sonication. The NP reaction mixture was allowed to react under sonication for 1 h and later collected into Eppendorf tubes for washing. The peptide-functionalized NPs were washed three times with nanopure water via centrifugation (10,000 × *g*, 5 min) and subsequently isolated magnetically with a DynaMag to remove unreacted peptide. The resulting peptide functionalized NPs were redispersed in nanopure water at a concentration of 5 mg/mL.

**Top-down RPLC-MS/MS Analysis of cTnI, cTnT, and TnC monomers.** For top-down RPLC-MS/MS protein sample preparation of the cardiac protein L, F, and NP-Pep E, Amicon Ultra Centrifugal filters (10 kDa, 0.5 mL) were pre-equilibrated with a cartridge volume of 0.2% formic acid (FA) in water and centrifuged at 14,000 × *g* for 5 min at 4 °C. After the filters were equilibrated, protein sample was introduced, and the proteins were desalted by repeated dilution with 0.2% FA in water to the cartridge filter volume, and concentrated by centrifugation at 14,000 × *g* for 5 min at 4 °C. This dilution and concentration process was repeated for five additional times to effectively buffer exchange proteins into MS-compatible mobile phase and to remove non-volatile salts and buffers. Buffer exchanged samples were then transferred to chilled LC-MS vials for reverse phase (RP)LC-MS/MS analysis. Top-down RPLC-MS/MS was carried out by either using an Acquity ultra-high pressure LC M-class system (Waters) coupled to a high-resolution maXis II quadrupole time-of-flight (QTOF) mass spectrometer (Bruker Daltonics) or by using a nanoAcquity ultra-high pressure LC system (Waters) coupled to a high-resolution Impact II QTOF mass spectrometer (Bruker Daltonics). 600 ng of total protein was injected onto a home-packed PLRP column (PLRP-S) (Agilent Technologies), 10-µm particle size, 500-µm inner diameter, 1,000 Å pore size using an organic gradient of 20 to 65% mobile phase B (mobile phase A: 0.2% FA in $H_2O$; mobile phase B: 0.2% FA in 50:50 acetonitrile/isopropanol) at a constant flow rate of 12 µL/min. For the maXis II QTOF, mass spectra were taken at a scan rate of 0.5 Hz over 200-3000 *m/z* range. Ion source parameters, the end plate offset, and capillary voltage were 500 and 4500 V, respectively. The nebulizer gas pressure ($N_2$), dry gas flow rate, and dry gas temperature were set to 0.5 bar, 4.0 L/min, and 200 °C, respectively. For ion transfer parameters, the funnel RF, octupole RF, quadrupole energy, low mass cutoff, and in-source collision-induced dissociation (isCID) energy were optimized to 400 V, 400 V, 5 V, 500 *m/z,* and 10 V, respectively. Additionally, collision Vpp, pre-pulse storage time, and collision cell transfer time were set to 2000 Vpp, 15 µs, and 100 µs, respectively. For the Impact II QTOF, mass spectra were taken at a scan rate of 0.5 Hz over 500-2000 *m/z* range. Ion source parameters, the end plate offset, and capillary voltage were 500 V and 4500 V, respectively. The nebulizer gas pressure ($N_2$), dry gas flow rate, and dry gas temperature were set to 0.5 bar, 4.0 L/min, and 200 °C, respectively. For ion transfer parameters, the funnel RF, hexapole RF, quadrupole energy, low mass cutoff, and in-source collision-induced dissociation (isCID) energy were optimized to 400 V, 400 V, 5 V, 500 *m/z,* and 10 V, respectively. Additionally, collision Vpp, pre-pulse storage time, and collision cell transfer time were set to 2000 Vpp, 15 µs, and 100 µs, respectively.

**Size exclusion chromatography (SEC) for online buffer exchange (OBE) and native MS analysis of the cTn(I-T-C) complex.** SEC experiments were performed using a NanoAcquity ultra-high pressure LC system (Waters) coupled to a high-resolution maXis II quadrupole time-of-flight mass spectrometer (Bruker Daltonics). 1 µg of total protein was injected onto a

PolyHYDROXYETHYL A column (PolyHEA) (PolyLC Inc), 2.1 mm internal diameter, 100 mm length, 5 µm particle size, and 200 Å pore size. Protein samples were separated isocratically with with 200 mM ammonium acetate solution at a flow rate of 28 µL/min for 10 min with a 'divert to waste' step programmed at 7 min. After 7 min, the SEC eluate consisted of non-volatile buffers and salts which were diverted from the electrospray ionization source to prevent fouling of the mass spectrometer. After every protein sample, the PolyHEA column was flushed with 1 column volume of HPLC $H_2O$, and re-equilibrated with 1 column volume of 200 mM ammonium acetate solution. Mass spectra were taken at a scan rate of 0.25 Hz over 500-8000 *m/z* range. For ion source parameters, the end plate offset, and capillary voltage were 500 V and 4500 V, respectively. The nebulizer gas pressure ($N_2$) was set to 1.5 bar, the dry gas flow rate set to 5.0 L/min, and the dry gas temperature set to 200°C. For ion transfer parameters, the funnel RF, octopol RF, quadrupole energy, low mass cutoff, and isCID were set to 400 V, 800 V, 4V, 500 *m/z*, and 150-200 V, respectively. Additionally, collision Vpp, pre-pulse storage time, and transfer time was optimized to 4000 Vpp, 45 µs, and 145 µs, respectively.

### Supplementary Tables

**Table S1. Available clinical data for non-failing donor hearts used.** Non-failing donor heart tissue have been coded to Donor#. Samples were obtained from the University of Wisconsin (UW)-Madison Organ and Tissue Donation. The sample code (assigned for this publication) and Biopsy ID are provided. Clinical data such as age, gender, cause of death, and medical history data are listed.

| Sample Code | Biopsy ID | Age | Gender | Cause of Death | Medical History |
|---|---|---|---|---|---|
| Donor 1 | YG53 | 65 | F | Head Trauma | Trivial mitral valve regurgitation, mild hypertension, smoking, anxiety |
| Donor 2 | YG54 | 62 | M | Head Trauma | No past history of heart disease, had hypothyroidism and high cholesterol |
| Donor 3 | YG66 | 43 | F | Overdose | No past history of heart disease or comorbidities, mild asthma |
| Donor 4 | YG57 | 64 | F | Stroke | Hypertension, hyperlipidemia, smoking, and mild coronary artery and aortic valve calcification |
| Donor 5 | YG59 | 56 | M | Brain Death | Emphezema, high cholesterol, bipolar, depression smoker. |

**Table S2. cTn proteoforms identified.** Proteoform, modified forms, theoretical most abundant mass, experimental most abundant mass, and mass error for the proteoforms we identified. Modified proteoforms were manually identified based on highly accurate intact mass measurements and MS/MS data together with the prior knowledge on identified cTn proteoforms from previous publications[1-4] or Uniprot. Abbreviations: cardiac troponin T (cTnT, T); cardiac troponin I (cTnI, I); troponin C (TnC, C); N-terminal acetylation (N-acetyl); phosphorylation (phospho, *p*); bis-phosphorylation (2x phospho, *pp*); methionine (Met).

| Proteoform | Modified Forms | Theo Mass (Da) | Expt Mass (Da) | Mass Error (ppm) |
|---|---|---|---|---|
| TnC | N-acetyl | 18443.58 | 18443.87 | 15.8 |
| TnC | N-acetyl, 1 Ca(II) | 18481.47 | 18481.52 | 2.9 |
| TnC | N-acetyl, 2 Ca(II) | 18519.46 | 18519.46 | 0.2 |
| TnC | N-acetyl, 3 Ca(II) | 18557.41 | 18557.42 | 0.3 |
| *p*cTnT [aa 1-286] | aa 1-286, Met removal, N-acetyl | 34452.31 | 34452.35 | 1.2 |
| cTnT | Met removal, N-acetyl | 34500.44 | 34500.41 | 0.9 |
| *p*cTnT | Met removal, N-acetyl, phospho | 34580.41 | 34580.34 | 2.0 |
| cTn(I-C) | TnC: N-actetyl, 2 Ca(II)<br>cTnI: Met-removal, N-acetyl | 42438.31 | 42438.32 | 0.2 |
| cTn(I-C) | TnC: N-acetyl, 3 Ca(II)<br>cTnI: Met-removal, N-acetyl | 42476.25 | 42476.27 | 0.5 |
| cTn(I-C) | TnC: N-actetyl, 2 Ca(II)<br>cTnI: Met-removal, N-acetyl, phospho | 42518.27 | 42518.32 | 1.2 |
| cTn(I-C) | TnC: N-actetyl, 3 Ca(II)<br>cTnI: Met-removal, N-acetyl, phospho | 42556.21 | 42556.26 | 1.1 |
| cTn(I-C) | TnC: N-actetyl, 2 Ca(II)<br>cTnI: Met-removal, N-acetyl, 2x phospho | 42598.23 | 42598.31 | 1.9 |
| cTn(I-C) | TnC: N-actetyl, 3 Ca(II)<br>cTnI: Met-removal, N-acetyl, 2x phospho | 42636.18 | 42636.17 | 0.3 |
| cTn(I-T-C) | TnC: N-actetyl, 2 Ca(II)<br>cTnI: Met-removal, N-acetyl, phospho<br>cTnT: Met-removal, N-acetyl, phospho | 77098.68 | 77098.73 | 0.8 |
| cTn(I-T-C) | TnC: N-actetyl, 3 Ca(II)<br>cTnI: Met-removal, N-acetyl, phospho<br>cTnT: Met-removal, N-acetyl, phospho | 77136.62 | 77136.68 | 0.7 |
| cTn(I-T-C) | TnC: N-actetyl, 2 Ca(II)<br>cTnI: Met-removal, N-acetyl, 2x phospho<br>cTnT: Met-removal, N-acetyl, phospho | 77178.64 | 77178.70 | 0.8 |
| cTn(I-T-C) | TnC: N-actetyl, 3 Ca(II)<br>cTnI: Met-removal, N-acetyl, 2x phospho<br>cTnT: Met-removal, N-acetyl, phospho | 77216.59 | 77216.68 | 1.2 |

**Table S3.** Summary of native TIMS-MS CCS (Å$^2$) values for TnC monomer, cTn(I-C) dimer, and cTn(I-T-C) heterotrimer complex. Theoretical CCS values were determined using the IMPACT[5] method.

| cTn | Theoretical CCS [Å$^2$] | Experimental CCS [Å$^2$] | Charge State [*z*+] |
|---|---|---|---|
| TnC | 2149 | 1853 | 8+ |
| TnC + 1Ca(II) | 2141 | 1849 | 8+ |
| TnC + 2Ca(II) | 2141 | 1829 | 8+ |
| TnC +3Ca(II) | 2131 | 1844 | 8+ |
| cTn (I-C) + 2Ca(II) | 3630 | 3623 | 15+ |
| cTn (I-C) + 3Ca(II) | 3637 | 3640 | 15+ |
| cTn(I-T-C) Complex | 4192 | 4574 | 19+ |

**Table S4.** Summary of native TIMS-MS CCS (Å$^2$) values for the cTn(I-T-C) complex incubated with 0, 25, 50, and 100 mM EGTA.

| EGTA Concentration (mM) | Charge state [*z*+] | Mobility [1/K0] | CCS [Å$^2$] |
|---|---|---|---|
| 0 | 20 | 1.16 | 4880.3 |
| 25 | 20 | 1.166 | 4905.2 |
| 50 | 20 | 1.174 | 4936.7 |
| 100 | 20 | 1.22 | 5132.1 |

**Supplementary Figures**

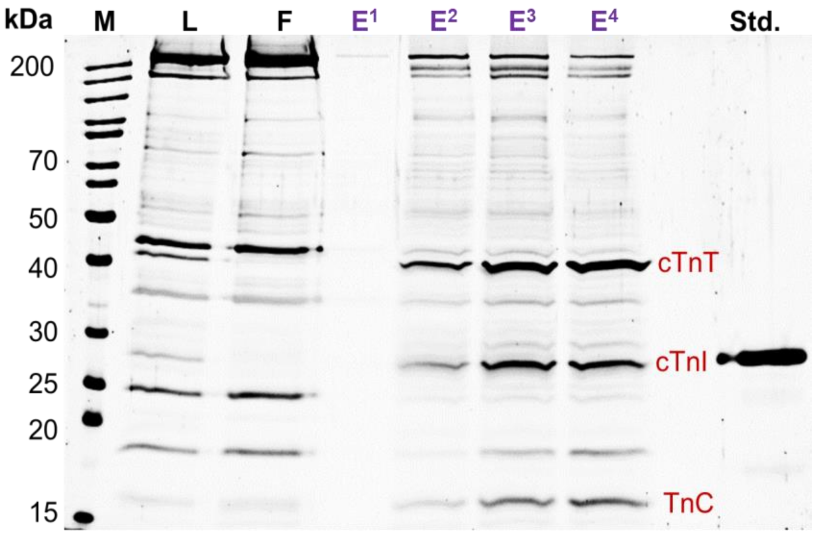


**Figure S1. Optimization of native elution buffer solution for cTn enrichment with NP-Pep.** SDS-PAGE visualizing sarcomeric proteins extracted from non-failing healthy donor heart tissue after the NP-Pep enrichment procedure. Equal amount (500 ng) of the loading mixture (L), flow-through (F), and elution mixture after enrichment (E) was loaded on the gel. $E^1$: 100 mM L-arginine, 100 mM imidazole, 50 mM L-glutamic acid (pH = 7.5); $E^2$: 250 mM L-arginine, 250 mM imidazole, 50 mM L-glutamic acid (pH = 7.5); $E^3$: 500 mM L-arginine, 500 mM imidazole, 50 mM L-glutamic acid (pH = 7.5); $E^4$: 750 mM L-arginine, 750 mM imidazole, 50 mM L-glutamic acid (pH = 7.5). Higher concentrations are required for more effective elution from NP-Pep, suggesting a mechanism of competitive elution. M. protein marker, Std. endogenous cTnI protein standard.

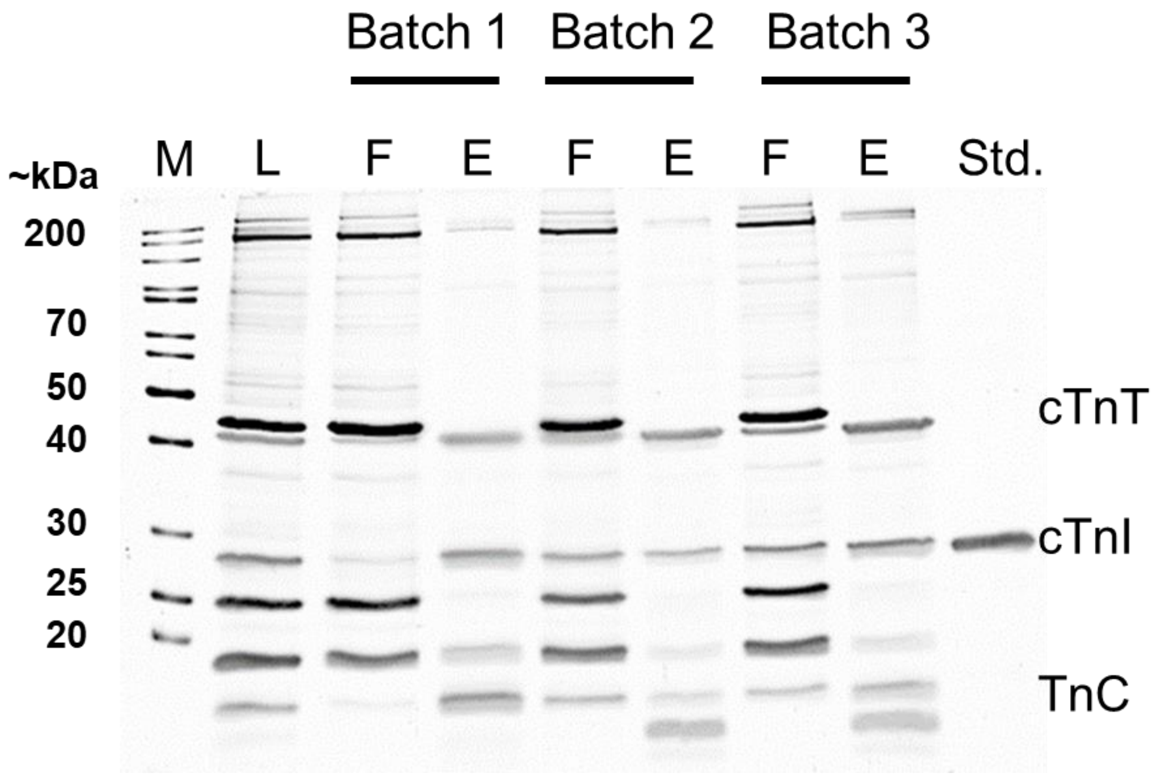


**Figure S2. High reproducibility of inter-batch enrichment by NP-Pep.** SDS-PAGE visualizing the cTn enrichment performance and demonstrating the high reproducibility obtained from three different synthesis batches of NP-Pep, using 750 mM L-Arg, 750 mM Imidazole, 50 mM L-Glu (pH = 7.5) as native elution buffer. Equal amount (500 ng) of the loading mixture (L), flow-through (F), and elution mixture after enrichment (E) was loaded on the gel. M. protein marker, Std. endogenous cTnI protein standard.

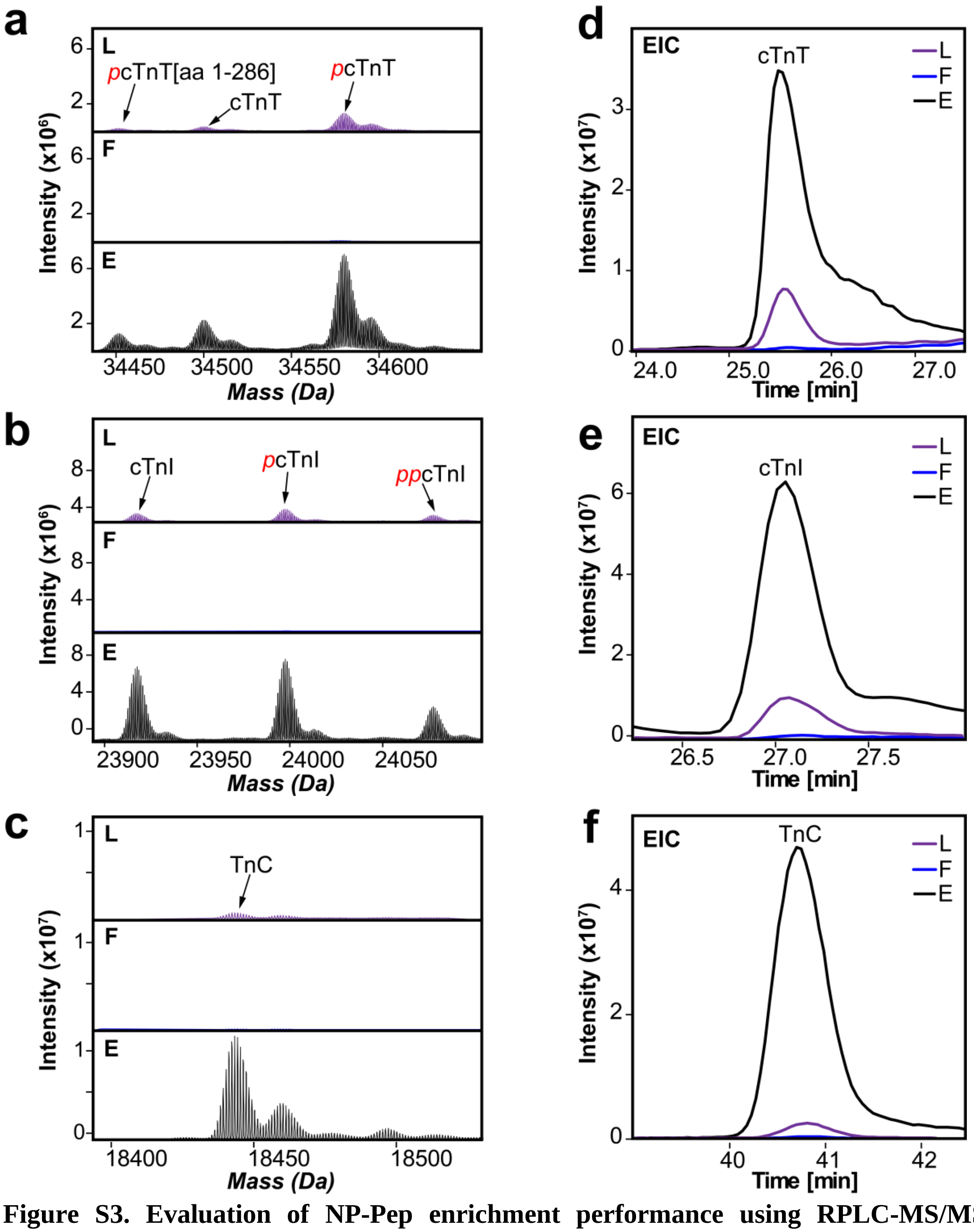


**Figure S3. Evaluation of NP-Pep enrichment performance using RPLC-MS/MS.** The deconvoluted top-down mass spectra of (a) cTnT, (b) cTnI, and (c) TnC proteoforms when loading mixture (L, purple), flow-through (F, blue), and elution mixture after native cTn enrichment (E, black) were equally loaded (600 ng) for online RPLC-MS/MS. Enrichment using NP-Pep demonstrates preservation of proteoforms while significantly enriching cTnT, cTnI, and TnC subunits compared to the initial L. Extracted ion chromatograms (EICs) of (d) cTnT, (e) cTnI, and (f) TnC for L, F, and E demonstrate successful enrichment of cTnT, cTnI, and TnC in E compared to the initial L.

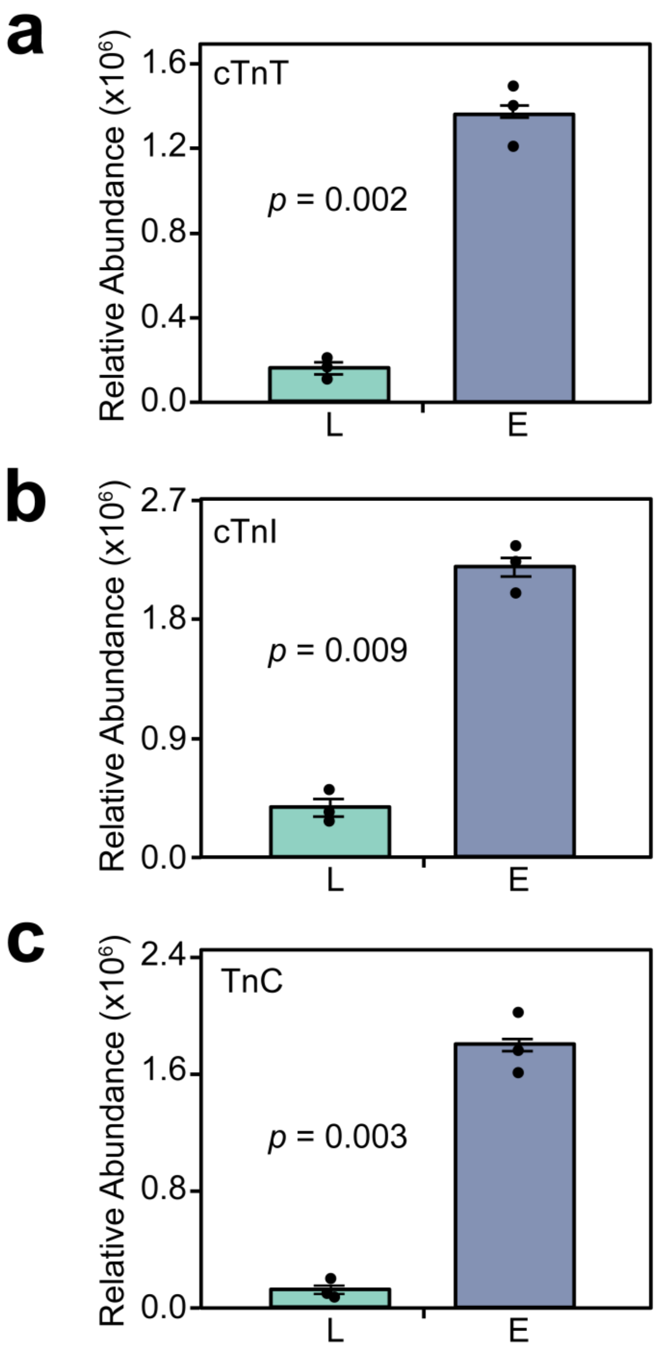


**Figure S4. Reproducibility of the NP-Pep enrichment capturing cTn subunits in human heart tissues using RPLC-MS/MS.** The relative abundance of cTn subunits was calculated using the deconvoluted top-down mass spectra corresponding to cTnT, cTnI, and TnC. cTnT (a), cTnI (b), and TnC (c) relative abundance data in the loading mixture (L) and elution mixture (E) are representative of $n = 3$ independent donor heart enrichments with error bars indicating standard error of the mean. Groups were considered significantly different by paired-students *t*-tests with $p < 0.01$.

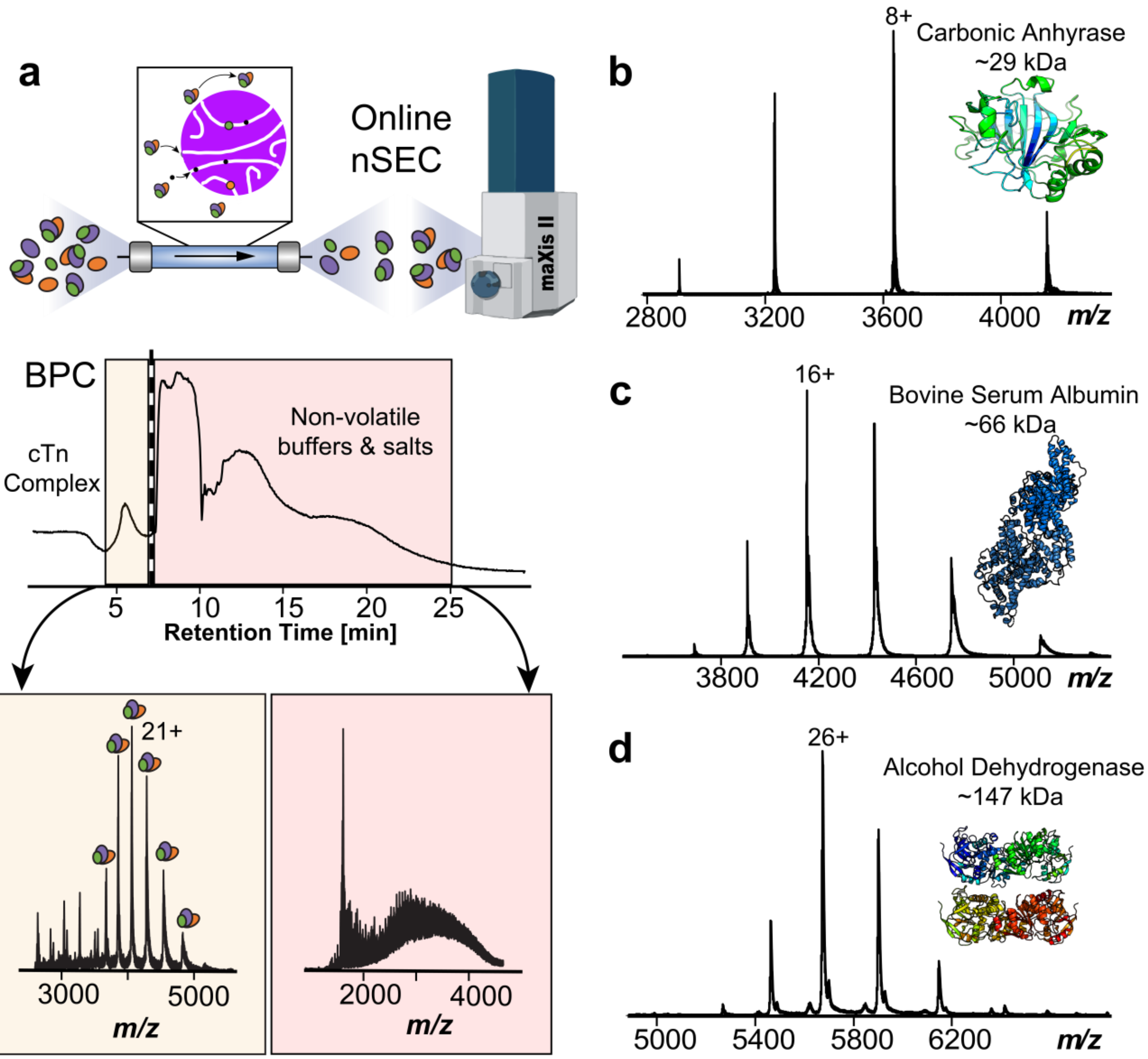


**Figure S5. Native size exclusion chromatography (SEC) for online buffer exchange (OBE) enables MS analysis of protein and protein complexes.** (a) Schematic representation of the native SEC-OBE workflow. Protein and protein complexes are separated and buffer exchanged into 200 mM ammonium acetate using a PolyHYDROXYETHYL A column (PolyHEA) column coupled to a Bruker maXis II quadrupole time-of-flight mass spectrometer for native MS data collection. Base peak chromatogram (BPC) of enriched cTn mixture, where the cTn complex (~77 kDa) elutes between 5-6 min. A 'divert to waste' step is programmed at 7 min to divert SEC eluate consisting of non-volatile buffer and salts from the electrospray ionization source. The native SEC-OBE MS method can also be applied to other standard protein and protein complexes including (b) carbonic anhydrase (~29 kDa, PDB: 12CA), (c) bovine serum albumin (~66 kDa, PDB: 6QS9), and (d) alcohol dehydrogenase tetramer (~144 kDa, PDB: 4W6Z).

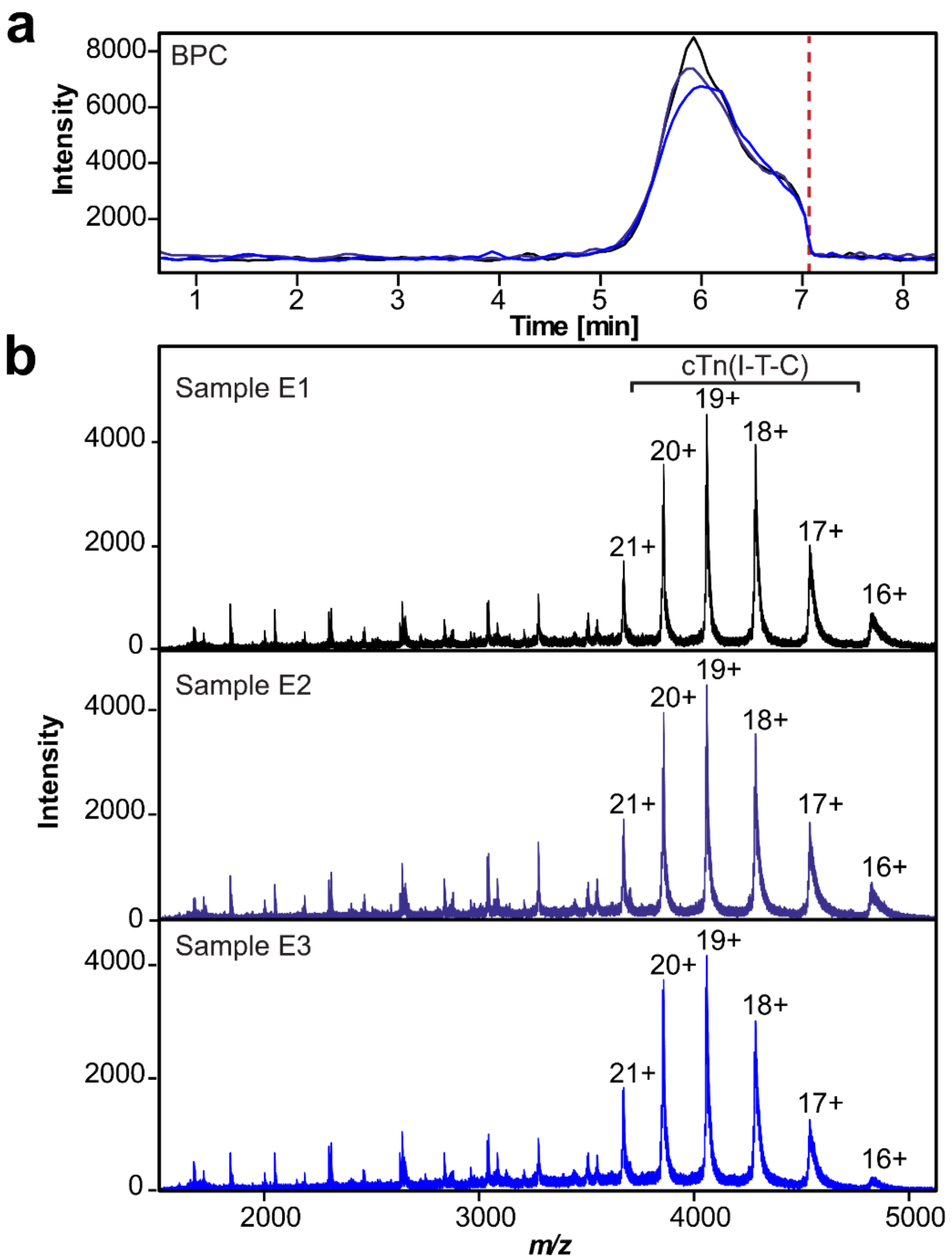


**Figure S6. Reproducibility of native cTn enrichment and SEC-OBE native MS analysis.** (a) Overlaid base peak chromatograms (BPCs) showing reproducibility of the intact cTn heterotrimer complex across three independent native enrichments using the same donor heart sample. Dashed red line is when the sample is diverted to waste to prevent non-volatile buffers and salts from fouling the electrospray ionization source of the mass spectrometer. (b) Native mass spectra of the three enrichment samples (E1, E2, and E3) demonstrating reproducible enrichment and SEC-OBE native MS analysis of the intact cTn heterotrimer complex ($z$ = 16-21+).

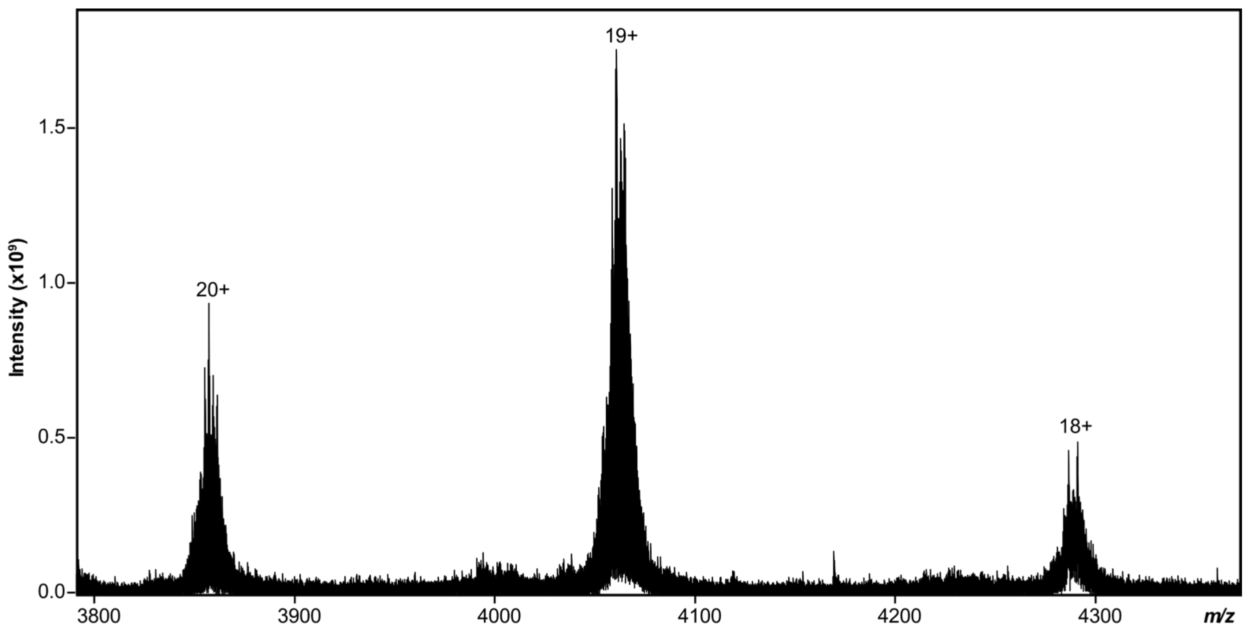


**Figure S7. Isotopically resolved native MS spectra of endogenous cTn heterotrimer complex (~77 kDa).** Native FTICR-MS mass spectrum of the heterotrimeric cTn complex directly enriched from human heart tissue. Charge states *z* = 18-20+ are shown.

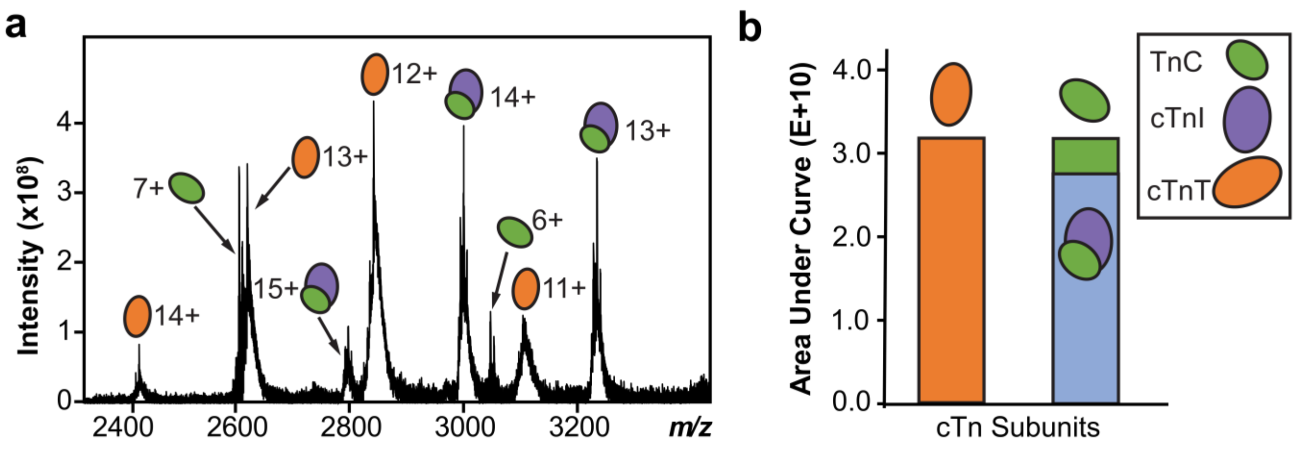


**Figure S8. Complex-up native MS analysis reveals cTn complex stoichiometry. (a)** Mass spectrum showing ejection of cTn subunits from the intact cTn(I-T-C) heterotrimer complex during complex-up analysis. **(b)** Bar graph showing the area under the curve (AUC) obtained from the three most abundant charge states of TnC monomer, cTnT monomer, cTn(I-C) dimer, and cTn(I-T-C) heterotrimer complex. There is an observed 1:1:1 correspondence between the molecular abundance of dissociated subunits.

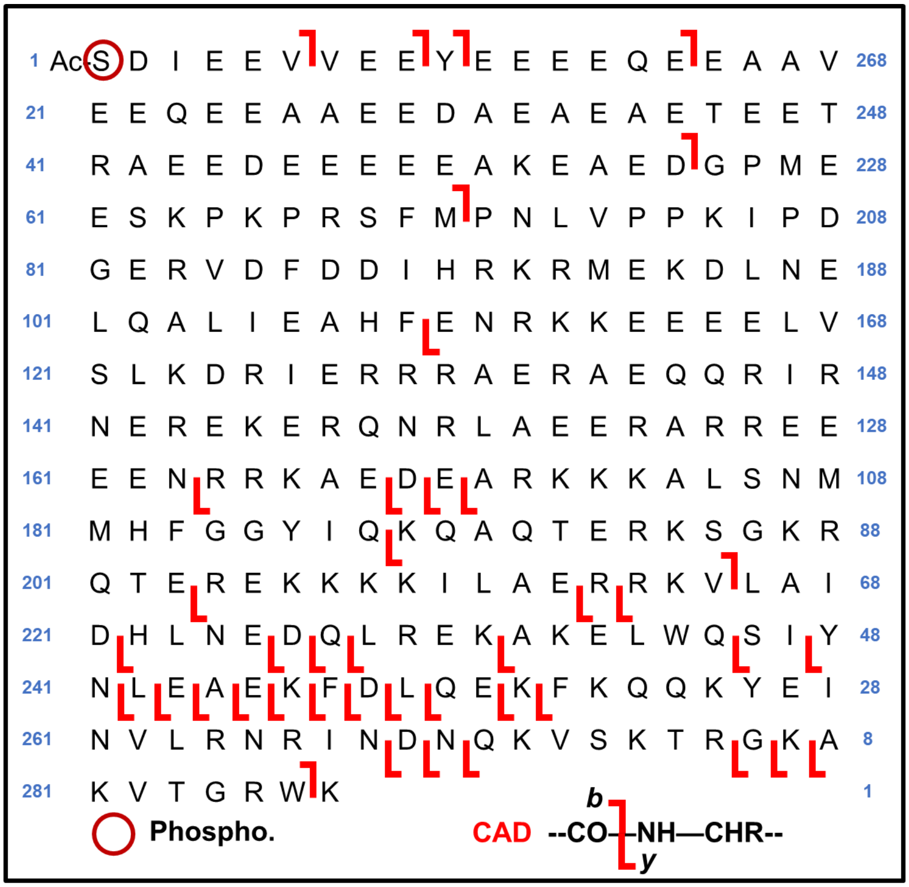


**Figure S9. Native top-down MS (nTDMS) fragmentation map of cTnT monomer.** Precursor ions were selected for collisionally activated dissociation (CAD) resulting in sequence informative b and y ions confirmed within a 20 ppm mass error tolerance. The sequence table with phosphorylation circled in red shows the characterization of cTnT monomer. The nTDMS data for cTnT generated 8 *b*-fragments and 33 *y*-fragments achieving 15% total bond cleavage.

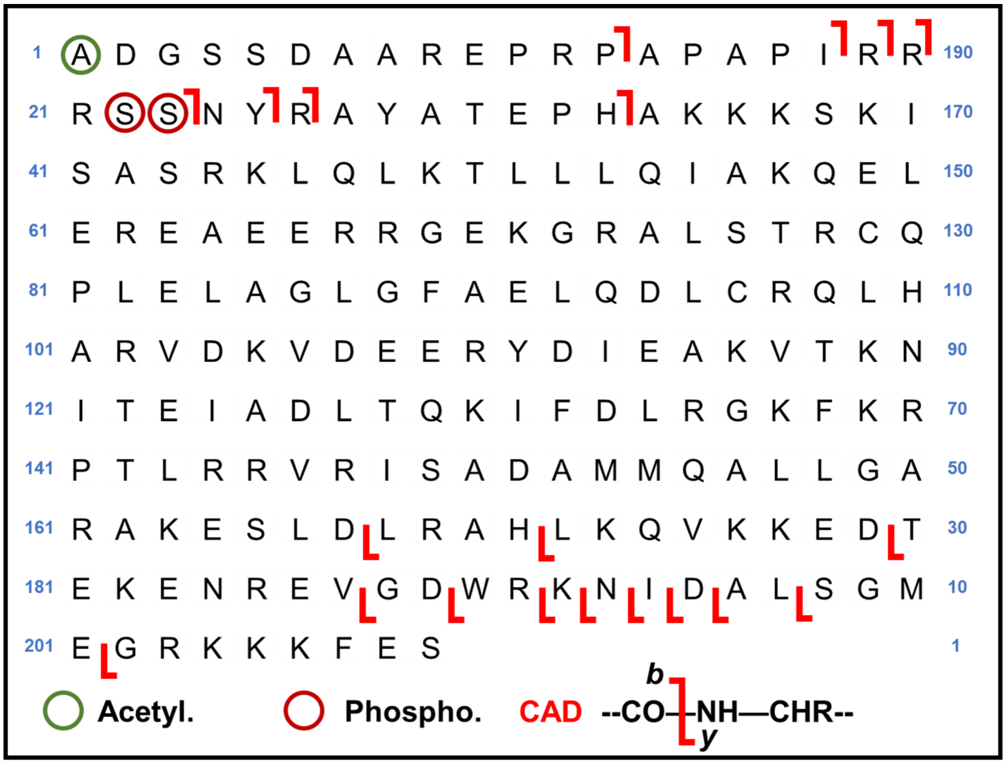


**Figure S10. Native top-down MS (nTDMS) fragmentation map of cTnI monomer.** Precursor ions were selected for collisionally activated dissociation (CAD) resulting in sequence informative b and y ions confirmed within a 20 ppm mass error tolerance. The sequence table with N-terminal acetylation circled in green and phosphorylation circled in red shows the characterization of cTnI monomer. The nTDMS data for cTnI generated 8 *b*-fragments, 12 *y*-fragments achieving 10% total bond cleavage.

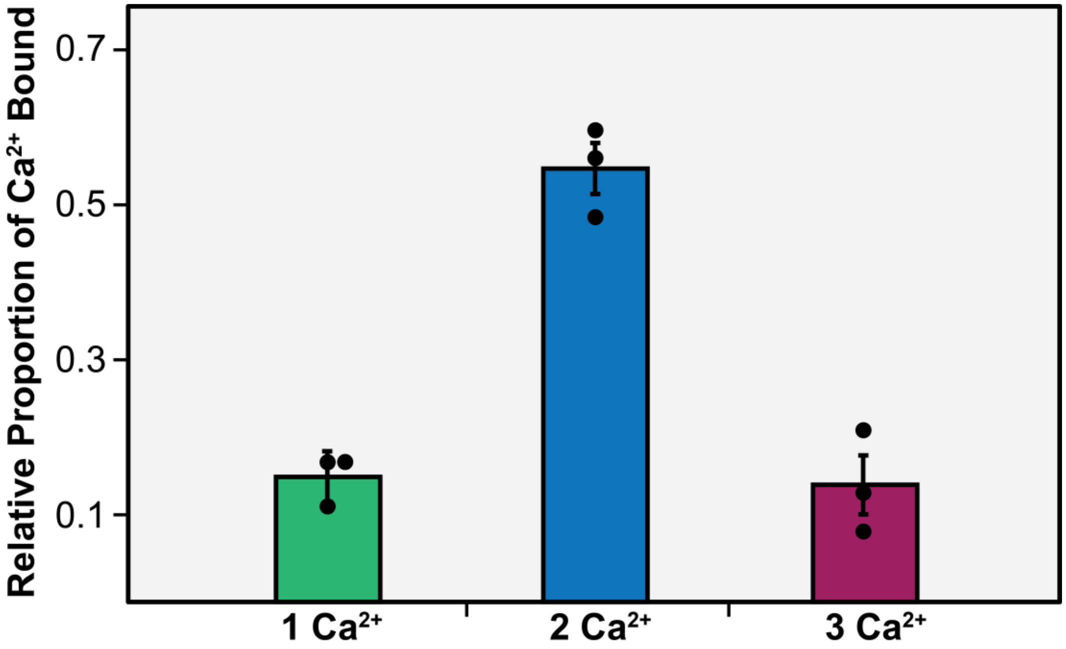


**Figure S11. Relative proportion of $Ca^{2+}$ bound in each TnC binding domain.** Relative $Ca^{2+}$ concentrations were quantified based on the ratio of the peak intensity of a $Ca^{2+}$-containing proteoform normalized to the summed peak intensities of all detected TnC proteoforms. Error bars indicate the mean ± standard error of the mean (S.E.M.). Data are representative of $n = 3$ independent experiments.

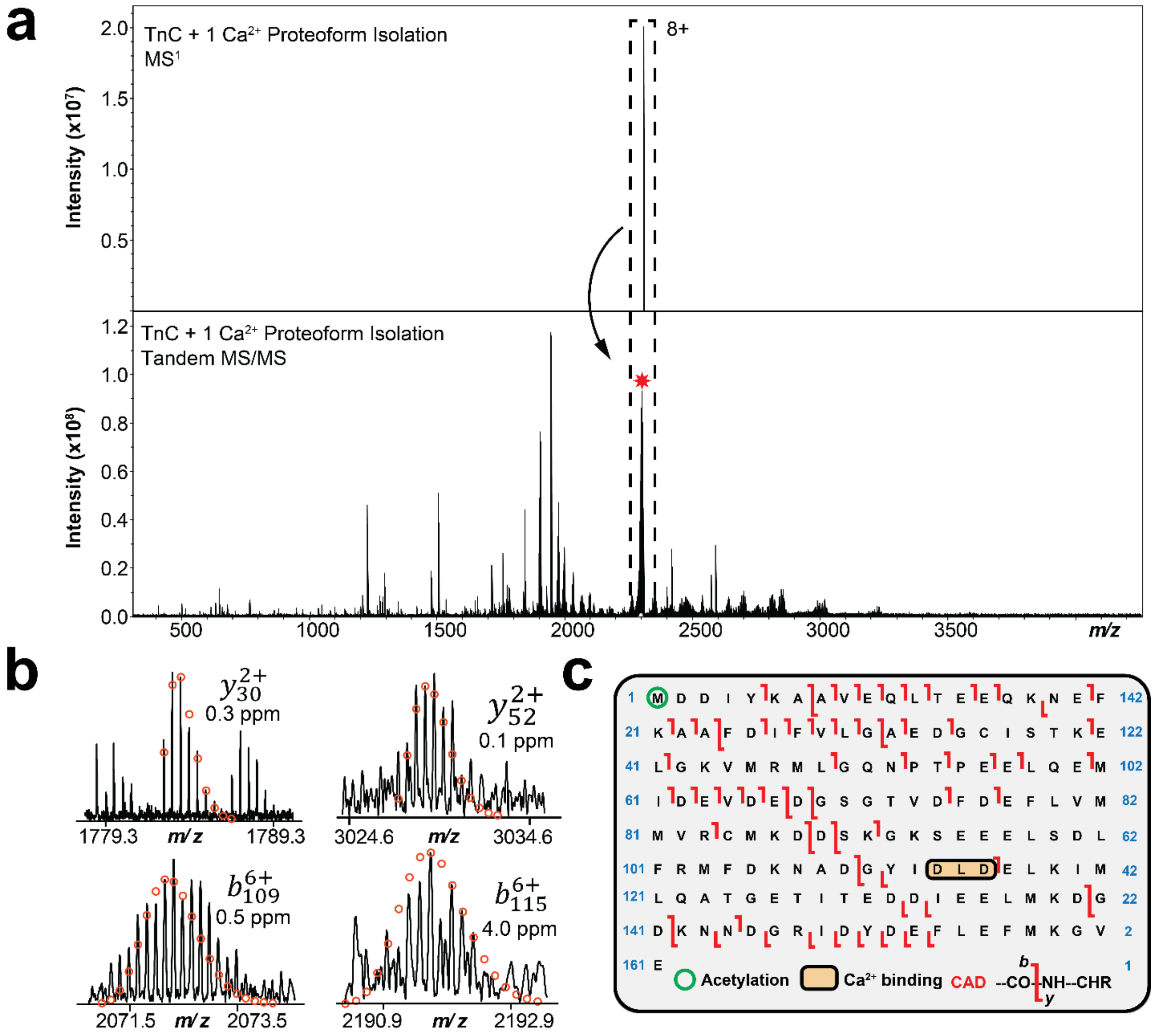


**Figure S12. Localization of calcium binding in TnC monomer domain III by native top-down MS (nTDMS).** (a) Isolation of native TnC monomer + 1 Ca(II) ($z$ = 8+, centered at 2312 $m/z$) from MS followed by nTDMS analysis of the isolated precursor. (b) Representative collionsally activated dissociation (CAD) fragment ions ($y_{30}^{2+}$, $y_{52}^{2+}$, $b_{109}^{6+}$, $b_{115}^{6+}$) obtained from the nTDMS analysis. Theoretical ion distributions are indicated by the red circles and mass accuracy errors are listed for each fragment ion. (c) Ca(II) localized to D113-D115 in TnC domain III.

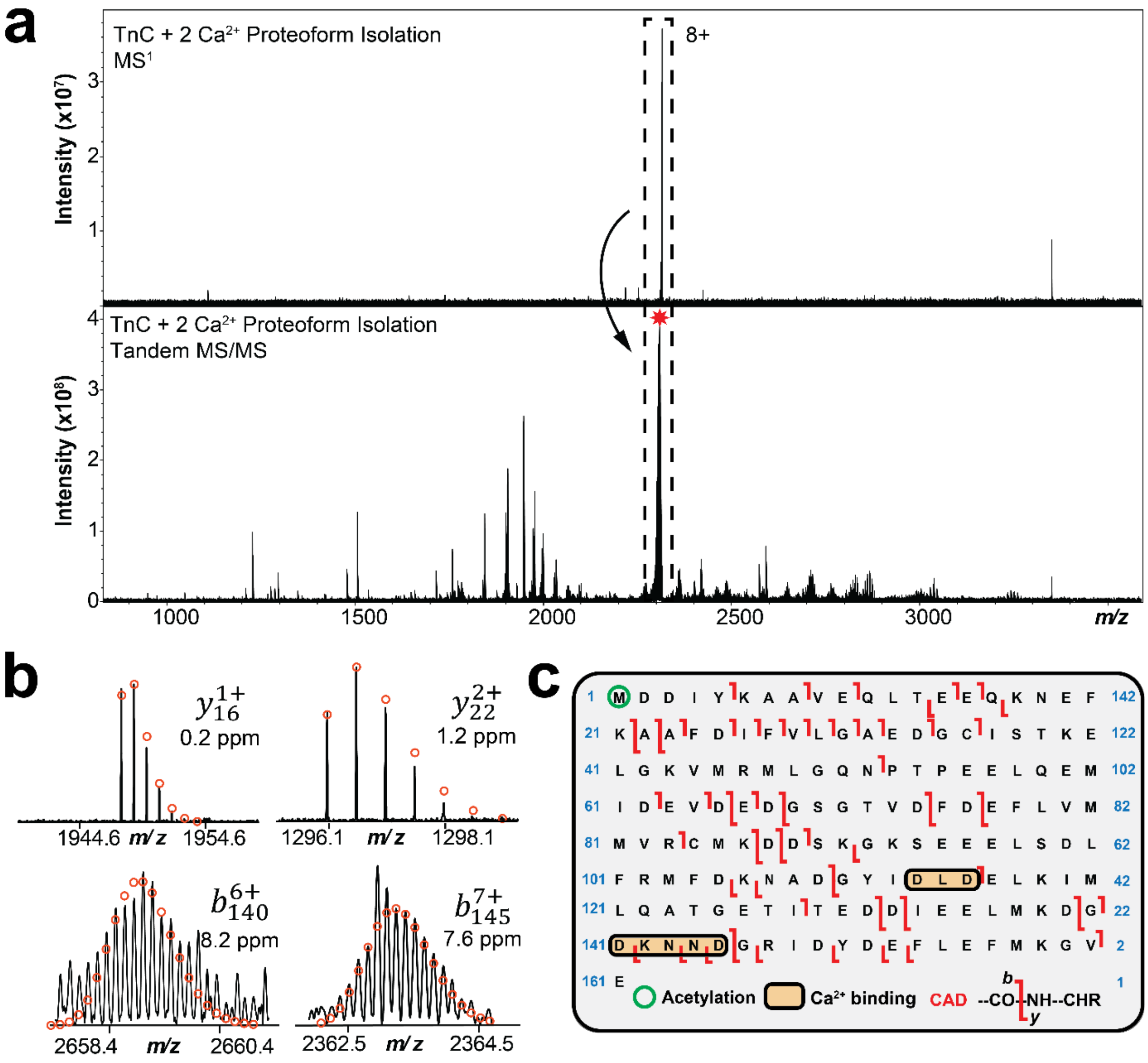


**Figure S13. Localization of calcium binding in TnC monomer domain IV by native top-down MS (nTDMS).** (a) Isolation of native TnC monomer + 2 Ca(II) ($z$ = 8+, centered at 2316 $m/z$) from MS followed by nTDMS analysis of the isolated precursor. (b) Representative collisionally activated dissociation (CAD) fragment ions ($y_{16}^{1+}$, $y_{22}^{2+}$, $y_{140}^{6+}$, $y_{145}^{7+}$) obtained from the nTDMS analysis. Theoretical ion distributions are indicated by the red circles and mass accuracy errors are listed for each fragment ion. (c) Ca(II) localized to D141-D145 in TnC domain IV.

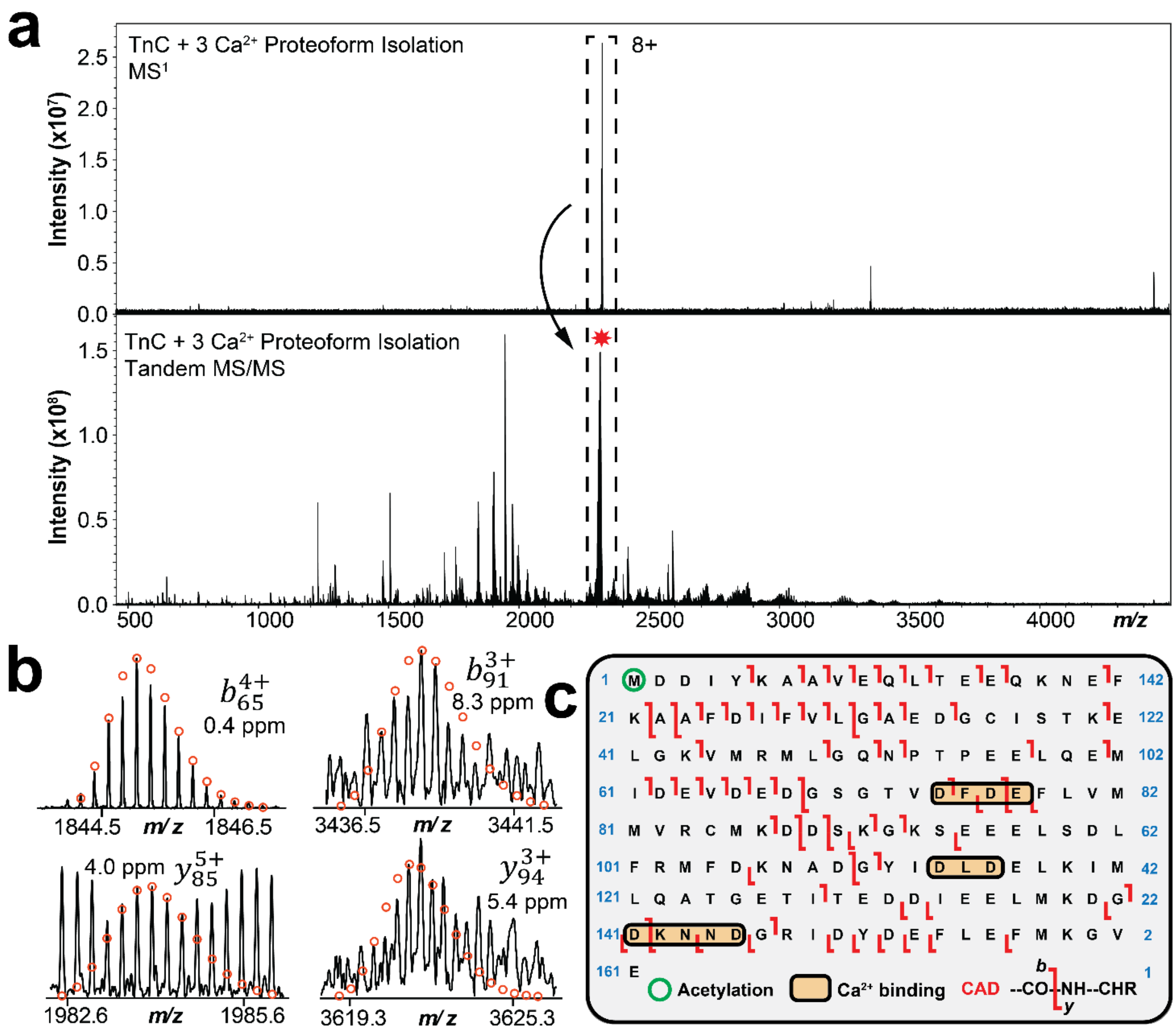


**Figure S14. Localization of calcium binding in TnC monomer domain II by native top-down MS (nTDMS).** (a) Isolation of native TnC monomer + 3 Ca(II) ($z$ = 8+, centered at 2321 $m/z$) from MS followed by nTDMS analysis of the isolated precursor. (b) Representative collionsally activated dissociation (CAD) fragment ions ($b_{65}^{4+}$, $b_{91}^{3+}$, $b_{85}^{5+}$, $b_{94}^{3+}$) obtained from the nTDMS analysis. Theoretical ion distributions are indicated by the red circles and mass accuracy errors are listed for each fragment ion. (c) Ca(II) localized to D73-E76 in TnC domain II.

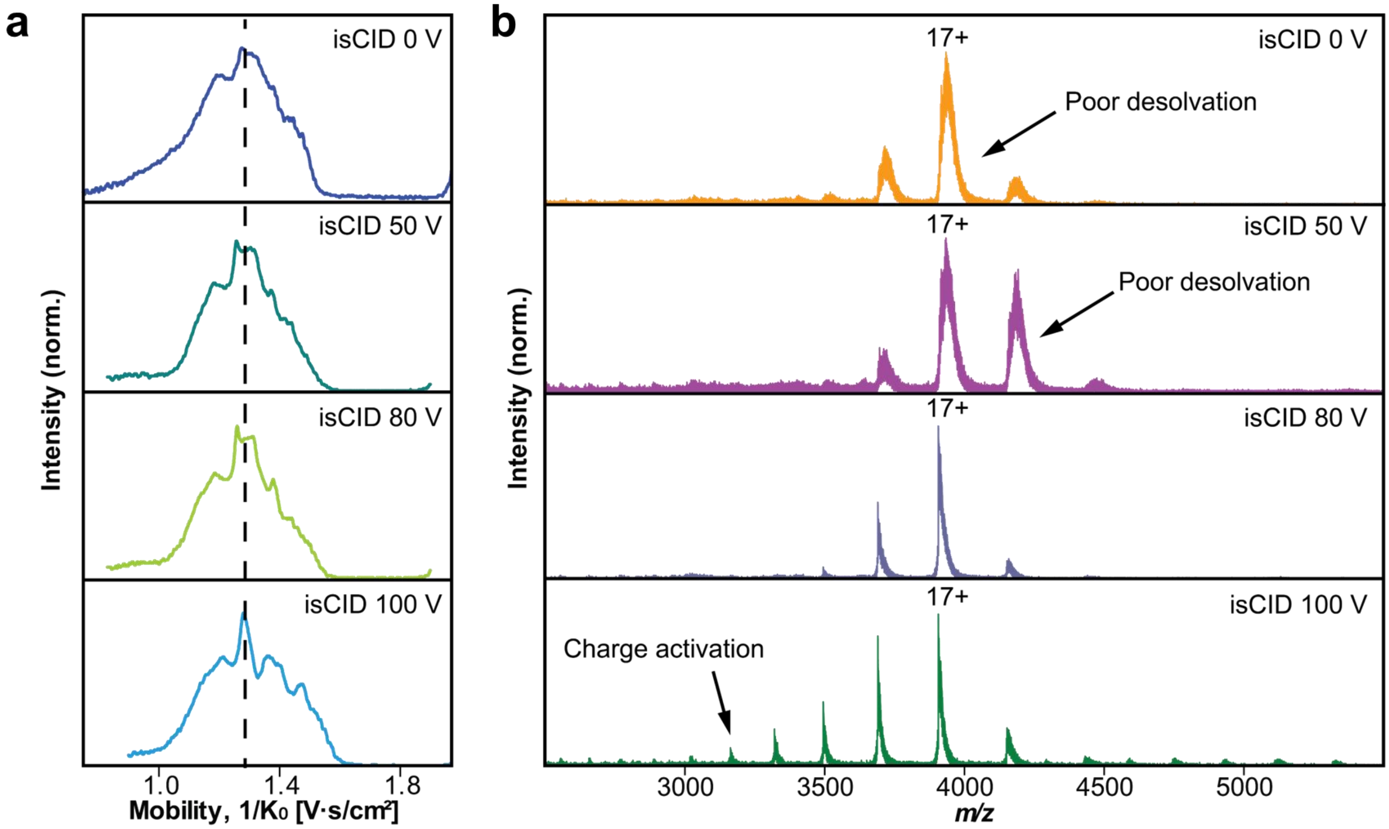


**Figure S15. Optimization of TIMS parameters for high resolution native MS analysis of bovine serum albumin (BSA).** (a) Normalized total ion mobility spectrum of native BSA showing distribution of $1/K_0$ (protein ion mobility) as a function of TIMS in-source collision-induced dissociation (isCID) (0-100V). (b) Normalized MS spectra of native BSA as a function of TIMS isCID. A TIMS isCID of 80 V yields the best desolvated gas-phase conformation of native BSA.

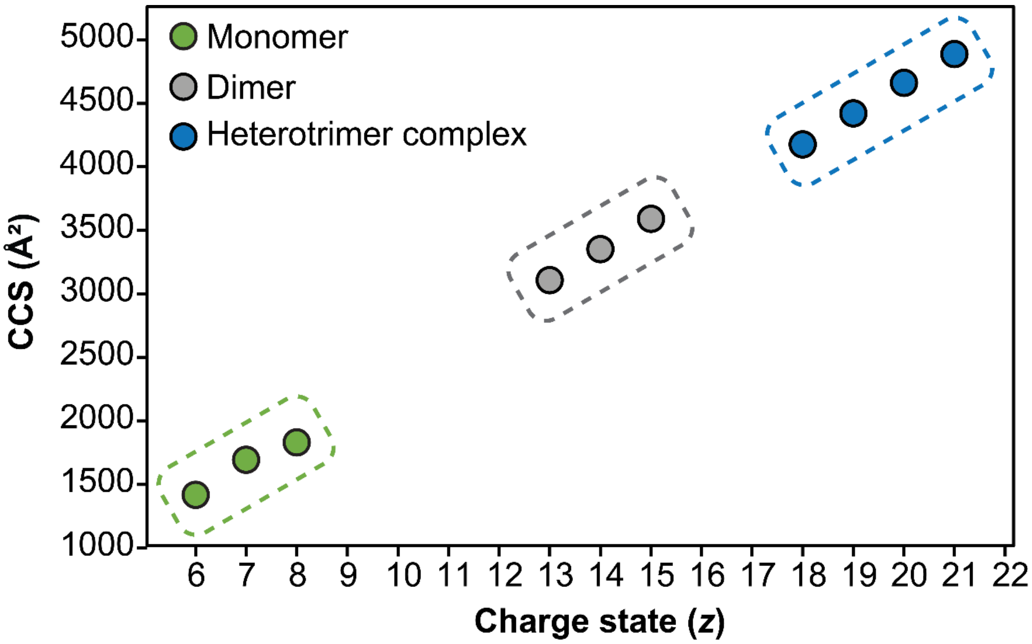


**Figure S16. Plot of CCS value as a function of charge state by native TIMS-MS.** Green circles indicate TnC monomer, grey circles indicate cTn(I-C) dimer charge states, and blue circles indicate cTn(I-T-C) heterotrimer complex charge states observed.
